# 6mer seed toxicity in viral microRNAs

**DOI:** 10.1101/838979

**Authors:** Andrea E. Murmann, Elizabeth T. Bartom, Matthew J. Schipma, Jacob Vilker, Siquan Chen, Marcus E. Peter

**Author notes:** Contact: Marcus Peter.

## Abstract

Micro(mi)RNAs are short double stranded noncoding RNAs (19-23nts) that regulate gene expression by suppressing mRNAs through RNA interference. Targeting is determined by the seed sequence (position 2-7/8) of the mature miRNA. A minimal G-rich seed of just 6 nucleotides is highly toxic to cells by targeting genes essential for cell survival. A screen of 215 miRNAs encoded by 17 human pathogenic viruses (v-miRNAs) now suggests that a number of v-miRNAs can kill cells through a G-rich 6mer sequence embedded in their seed. Specifically, we demonstrate that miR-K12-6-5p, an oncoviral mimic of the tumor suppressive miR-15/16 family encoded by human Kaposi’s sarcoma-associated herpes virus, harbors a noncanonical toxic 6mer seed (position 3-8) and that v-miRNAs are more likely than cellular miRNAs to utilize a noncanonical 6mer seed. Our data suggest that during evolution viruses evolved to use 6mer seed toxicity to kill cells.

## INTRODUCTION

We recently identified a code embedded in the genome that is based on RNA interference (RNAi) and allows for the effective killing of cells (Gao et al., 2018). RNAi is a form of post-transcriptional regulation exerted by 19-23 nt long double stranded RNAs that negatively regulate gene expression at the mRNA level. For a micro(mi)RNA, the RNAi pathway begins in the nucleus with transcription of a primary miRNA precursor (pri-miRNA) (Lee et al., 2004). Pri-miRNAs are first processed by the Drosha/DGCR8 microprocessor complex into pre-miRNAs (Han et al., 2004), which are then exported from the nucleus to the cytoplasm (Yi et al., 2003). Once in the cytoplasm, Dicer processes them further (Bernstein et al., 2001; Hutvagner et al., 2001) and these mature dsRNA duplexes are then loaded into Argonaute (Ago) proteins to form the RNA-induced silencing complex (RISC) (Wang et al., 2008). Translational silencing by miRNAs can be initiated with as little as six nucleotide base-pairing between a guide RNA’s seed sequence (position 2 to 7) and fully complementary seed matches in the target RNA (Lai, 2002; Lewis et al., 2003). This seed-based targeting most often occurs in the 3’ untranslated region (UTR) of a target mRNA (Baek et al., 2008; Selbach et al., 2008).

While effective targeting by many miRNAs requires a seed of at least 7 nucleotides and also involves pairing beyond the seed region (Broughton et al., 2016), we previously discovered a group of short interfering (si) and short hairpin (sh)RNAs that kill all cancer cells through miRNA-like minimal targeting between the 6mer seed and hundreds of essential genes critical for cell survival (Putzbach et al., 2017). The downregulation of essential survival genes was likely the cause of cell death rather than an effect because it preceded the initiation of cell death by three days and targeting the most highly downregulated survival genes directly using very low concentrations of SmartPool siRNAs against these genes also killed the cells (Putzbach et al., 2017). We called this form of cell death DISE, for death induced by survival gene elimination (Putzbach et al., 2018a). Based on the observation that only six nucleotides of complementarity between the si/shRNA and the 3’ UTR of mRNAs was required for targeting, we screened all 4096 possible six nucleotide permutations in a neutral 19mer siRNA backbone to determine which seed sequences were the most toxic to cells. This screen was performed in two human and two murine cancer cell lines derived from lung, liver, and ovarian cancer (Gao et al., 2018). Regardless of the species or tissue of origin, the most toxic seed sequences showed an enrichment for Gs in the seed sequence, with an asymmetrical preference for Gs at the 5’ end of the seed. Complementary C-rich seed matches to the most toxic seeds were identified predominantly located at the beginning of the 3’ UTR of many essential survival genes (Gao et al., 2018). By analyzing all human miRNAs we found that certain tumor suppressive miRNAs contain G-rich 6mer sequences in the first 6 positions of their seed (e.g. miR-34a-5p) and can kill cells through what we have called "6mer seed toxicity". In fact, our data suggest that 6mer seed toxicity shaped the miRNA repertoire over at least the last 800 million years. miRNAs are generated from a hairpin that has two arms. Usually only one arm of a mammalian miRNA is expressed, the other one is degraded. Almost all of the highly conserved and therefore older miRNAs do not contain G-rich 6mer sequences in the seed region of their expressed arm. This selection likely occurs to avoid the inclusion of miRNA seed matches in the 3’ untranslated region (3’UTR) of housekeeping genes, an observation that was described previously (Stark et al., 2005). In contrast, many younger miRNAs (<10 million years) and a small number of very old tumor suppressive miRNAs carry a highly toxic 6mer seed in their predominant arm (Gao et al., 2018).

Viruses have coevolved with all cellular functions by either adapting to or by hijacking them for their own life cycle (Koonin et al., 2015). We therefore hypothesized that many viruses either avoid triggering cell death (e.g. by being AU-rich) or utilize this mechanism to affect cell fate (e.g. through viral miRNAs (v-miRNAs) carrying toxic 6mer seeds). Recently, the Kaposi’s sarcoma-associated herpes virus (KSHV) miRNA, miR-K12-6-5p, was identified as an oncoviral mimic of the human tumor suppressive miRNA family miR-15/16 (Morrison et al., 2019). KSHV is a human tumor virus that causes Kaposi’s sarcoma and primary effusion lymphoma, as well as the B cell proliferative disorder multicentric Castleman’s disease (Cesarman et al., 1995; Nador et al., 1996; Soulier et al., 1995). KSHV shows extensive “molecular piracy” of cellular factors and pathways (Moore and Chang, 1998) and of miRNAs. The virus encodes 20 miRNAs that are part of the viral latency program (Cai et al., 2005; Grundhoff et al., 2006; Pfeffer et al., 2005; Samols et al., 2005). kshv-miR-K11 shares a canonical 2 to 8 7mer seed with miR-155 and many of the same targets as miR-155 (Gottwein et al., 2007; Skalsky et al., 2007). In addition, the viral miR-K10a miRNA shares at least a subset of its targets with cellular miR-142-3p miRNA (Gottwein et al., 2007). Finally, miR-K12-6-5p was identified as a potential tumor suppressive miRNA mimicking miR-15/16 and has been posited to allow KSHV to buffer viral oncogene expression and avoid severe pathogenesis in the healthy host.

We now provide evidence that both miR-15/16 miRNAs and miR-K12-6-5p are toxic to cells in part based on the first 6 nucleotides of their seed sequences. In miR-K12-6-5p the toxic 6mer seed is offset by one nucleotide, starting at position three of the mature miRNA. To determine whether this activity is found throughout the space of all miRNAs that are encoded by viruses pathogenic to humans, we performed a toxicity screen of 215 v-miRNAs coded for by 17 human viruses and compared the toxicity and the targeting to a matching set of siRNAs carrying the same 6mer seed in a neutral backbone. We provide evidence that many v-miRNAs have a seed that contains a 6mer sequence that is toxic through G-richness and provide evidence that noncanonical 6mer seeds (pos. 3-8) contribute to this activity. The data not only validate the 6mer seed toxicity concept but also suggest that certain viruses evolved to use this mechanism to kill cells.

## RESULTS

### The miR-15a/b-5p/16-5p family kills cells in part through 6mer seed toxicity

We recently provided evidence that the master tumor suppressor miR-34a-5p kills cancer cells in large part through a G-rich 6mer sequence at the beginning of its seed (Gao et al., 2018). Transfecting cells with full length miR-34a-5p or just the miR-34a-5p 6mer seed in a neutral siRNA backbone was equally toxic to cells, resulting in almost 80% overlap in downregulated genes containing the same number of essential survival genes, and the downregulated genes were similarly enriched in genes carrying a miR-34a-5p 6mer seed match in their 3’UTR. The 6mer seed toxicity screen also identified a toxic 6mer seed that is shared by another established tumor suppressive miRNA family, the miR-15a/b-5p/miR-16-5p family (miR-15/16-5p for simplicity), which were the first miRNAs identified as tumor suppressors (Calin et al., 2002). Consistent with its lower predicted seed toxicity, it inhibited cell growth of HeyA8 cells less aggressively than miR-34a-5p (Gao et al., 2018). We set out to test how much of its toxicity stemmed from its embedded 6mer seed. By comparing the sequence of miR-15a-5p or miR-16-5p with that of the corresponding 6mer seed in the backbone used in the screen, we noticed that miR-15a-5p shared 10 nucleotides and miR-16-5p 8 nucleotides of extended seed identity with the oligonucleotide scaffold (Figure 1A, top). To test the contribution of only the first 6 nucleotides of the seed to the activity of the miRNAs, we modified the scaffold in the four positions that follow the 6mer seed by swapping the two complementary nucleotides to maintain the nucleotide composition in the duplex while eliminating the sequence identity with miR-15a-5p/16-5p (Figure 1A, bottom). When transfected into HeyA8 cells, the two miRNAs, the 6mer seed embedded in the original backbone (miR-15a-5p^10Seed^/16-5p^8Seed^) and the 6mer seed of these two miRNAs embedded in the modified backbone (miR-15a-5p/16-5p^6Seed^) all strongly reduced cell viability (Figure 1B). To determine how many genes were co-downregulated by the three reagents, we performed RNA-Seq analysis of the cells 48 hours after transfection. Consistent with the fact that miR-15a-5p and miR-16-5p share an 8mer seed sequence >78% of genes that were significantly downregulated (>1.5 fold, adj. p < 0.05) in cells transfected with miR-15a-5p were also downregulated in cells treated with miR-16-5p (Figure 1C). To determine the contribution of the first 6 nucleotides of the seed of each miRNA to its toxicity, we compared the downregulated genes in cells treated with either miR-15a-5p or miR-16-5p to the downregulated genes in cells treated with the siRNA sharing only the same 6mer seed. We found that 61% and >55%, respectively, of the genes that were downregulated in cells transfected with only the 6mer seed were also downregulated in the cells treated with the full miRNAs (Figure 1D, E). A gene ontology analysis revealed that the genes that were most downregulated grouped into GO terms that are consistent with cells dying from 6mer seed toxicity/DISE including mitosis, cell division, cell cycle regulation, and DNA replication (Gao et al., 2018; Putzbach et al., 2017; Putzbach et al., 2018b) (Figure 1D, E). A gene set enrichment analysis confirmed that in all cases essential survival genes but not a control set of nonsurvival genes (as previously defined (Putzbach et al., 2017)) were downregulated with similar significance and enrichment scores (Figure 1F). Finally, a Sylamer analysis to identify the seed matches most enriched in the 3’UTR of the downregulated genes demonstrated that in all three cases the same 6mer seed match was most highly enriched (Figure 1G). This analysis suggests a significant contribution of the first 6 nucleotides of the miRNA seed to the toxicity of the miR-15/16-5p family supporting our interpretation that a number of tumor suppressive miRNAs are toxic to cancer cells through 6mer seed toxicity.

**Figure 1.**
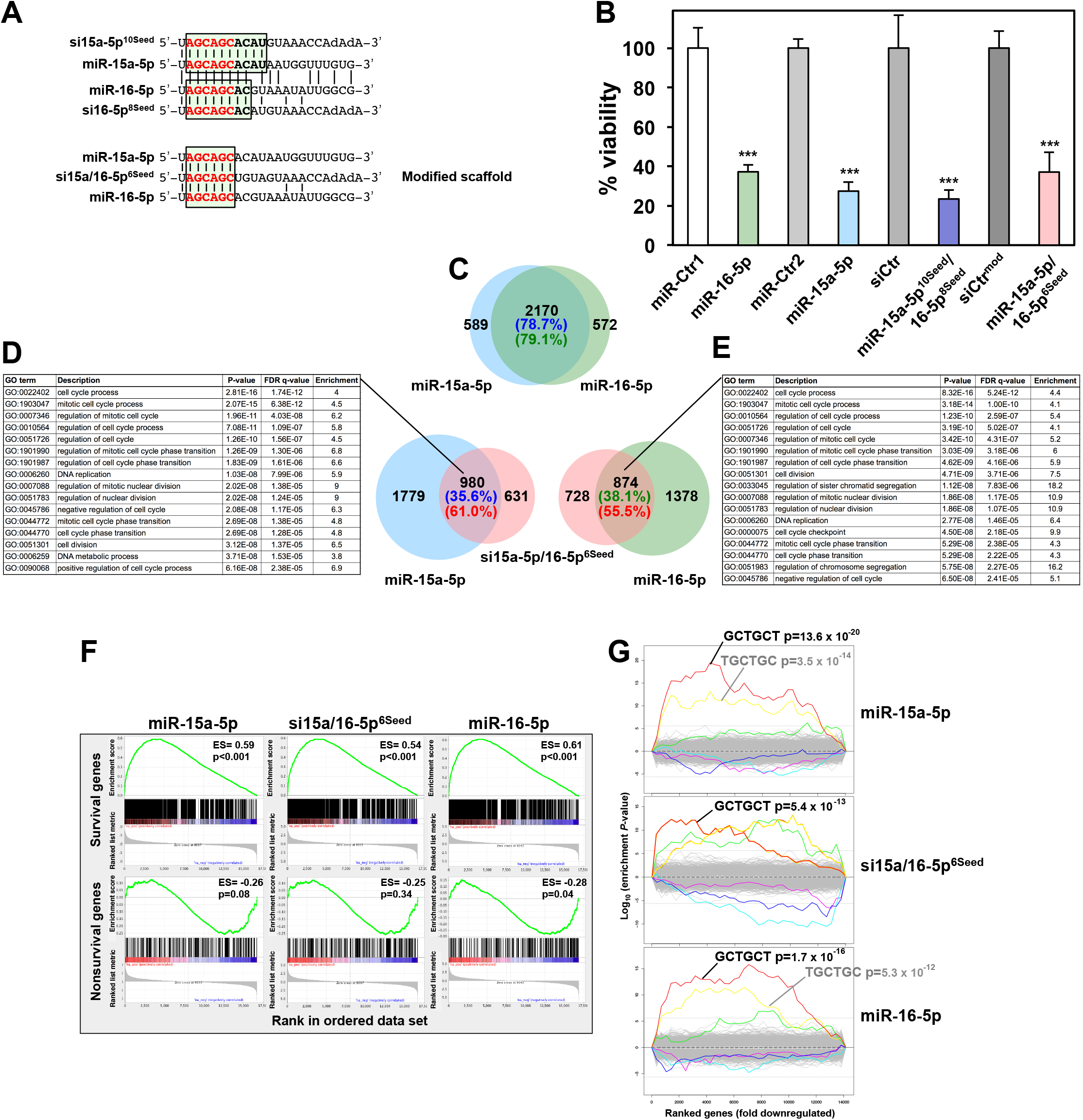
6mer seed toxicity in the tumor suppressive miR-15/16 miRNA family. (A) *Top:* The sequences of hsa-miR-15a-5p and hsa-miR-16-5p each aligned with the AGCAGC 6mer seed siRNA used in the previous screen with the extended seed identity highlighted (10 nucleotides for miR-15a-5p and 8 for miR-16-5p). The 6mer seed sequence is shown in red. *Bottom:* Alignment of miR-15a-5p and miR-16-5p with the AGCAGC 6mer seed siRNA modified to limit seed identity to 6 nucleotides. (B) Viability of HeyA8 cells 96 hours after transfection with 10 nM of either the indicated tumor-suppressive miRNAs or two miRNA non-targeting controls. Data are representative of two independent experiments. Each data point represents mean + SD of 5 replicates. *** p<0.0001 determined by Student’s T-test each compared to the respective control. (C) Overlap of RNAs detected by RNA-Seq downregulated in HeyA8 cells (>1.5-fold) 48 h after transfection with either miR-15-5p or miR-16-5p when compared to non-targeting miR-Ctr. (D) *Right:* Overlap of RNAs detected by RNA-Seq downregulated in HeyA8 cells (>1.5-fold) 48 h after transfection with either miR-15-5p or si15a/16-5p^6Seed^ compared to non-targeting miR-Ctr and modified siNT1, respectively. Left: Results of a GOrilla gene ontology analysis of the genes downregulated in cells transfected with miR-15a-5p or si15a/16-5p^6Seed^ (significance of enrichment <10−8). (E) *Left:* Overlap of RNAs detected by RNA-Seq downregulated in HeyA8 cells (>1.5-fold) 48 h after transfection with either miR-16-5p or si15a/16-5p^6Seed^ when compared to non-targeting miR-Ctr and modified siNT1, respectively. *Right:* Results of a GOrilla gene ontology analysis of the genes downregulated in cells transfected with miR-16-5p or si15a/16-5p^6Seed^ (significance of enrichment <10−8). (F) Gene set enrichment analysis for a group of 1846 survival genes (top three panels) and 416 non-survival genes (bottom three panels) after transfecting HeyA8 cells with either miR-15a-5p, miR-16-5p, or si15a/16-5p^6Seed^. mod-siNT1 and a non-targeting miR-Ctr served as controls, respectively. p-values indicate the significance of enrichment and the enrichment score (ES) is given. (G) Sylamer plots for the list of 3′UTRs of mRNAs in cells treated with either miR-15a-5p, miR-16-5p, or si15a/16-5p^6Seed^ sorted from downregulated to upregulated. For each plot the three most highly enriched ones are labeled in red, yellow and green, respectively. Bonferroni-adjusted p-values are shown.

### kshv-miR-K12-6-5p kills cells in part through 6mer seed toxicity

Given the apparent conservation of the 6mer seed toxicity mechanism during evolution we wondered whether human viruses would code for miRNAs that might be toxic to cells at least in part through this mechanism. There are currently 216 mature v-miRNAs coded by human pathogenic viruses listed in miRBase v.22.1 (**Table S1**). We searched this list of v-miRNAs for a miRNA that contains a position 2-7 seed that we determined to be toxic in our screen of 4096 6mer seeds and that was part of a major tumor suppressive miRNA family. While there was no v-miRNA with the same 6mer seed as miR-34a-5p, we found a v-miRNA that shared the same 2-7 nucleotides present in the predominant 5p arm of the miR-15/16-5p miRNA family. kshv-miR-K12-6-5p (miR-K12-6-5p) contains a GC-rich trinucleotide repeat seed sequence that is found in three miRNA families with ample published evidence for having tumor suppressive activities: miR-15/16-5p (Bonci et al., 2008; Calin et al., 2008; Calin et al., 2002; Han et al., 2017; Hu et al., 2018; Huang et al., 2015; Ke et al., 2013; Klein et al., 2010; Lovat et al., 2015; Lovat et al., 2018; Maximov et al., 2019; Pekarsky and Croce, 2015; Rahman et al., 2014; Yang et al., 2014), miR-214-3p (Cagle et al., 2019; Fan et al., 2017; Huang et al., 2014; Liu et al., 2016; Long et al., 2015; Pang et al., 2018; Phatak et al., 2016; Wang et al., 2012; Yang et al., 2015; Yang et al., 2018b; Zhang et al., 2017), and miR-103/107-3p (Datta et al., 2012; Feng et al., 2012; Fu et al., 2019; Gao et al., 2017; Piao et al., 2012; Sharma et al., 2017; Song et al., 2015; Yamakuchi et al., 2010; Yang et al., 2018a; Zhang et al., 2019) (Figure 2A). miR-K12-6-5p shares an 8mer seed sequence with miR-15a-5p, miR-15b-5p, and miR-16-5p, and a 6, 7 or 8mer seed with other members of this family (Figure 2A). However, the seed identity with miR-15/16-5p only occurs when the seed is counted from position 3 of the guide strand. A canonical 6mer seed identity can be found with the predominant arm in miRNAs related to miR-214 and a seed similarity with the predominant arm of the miR-103/107 family (miR-103a-3p/107) only exists when the seed is further offset so that it starts at position 4 in the v-miRNA.

**Figure 2.**
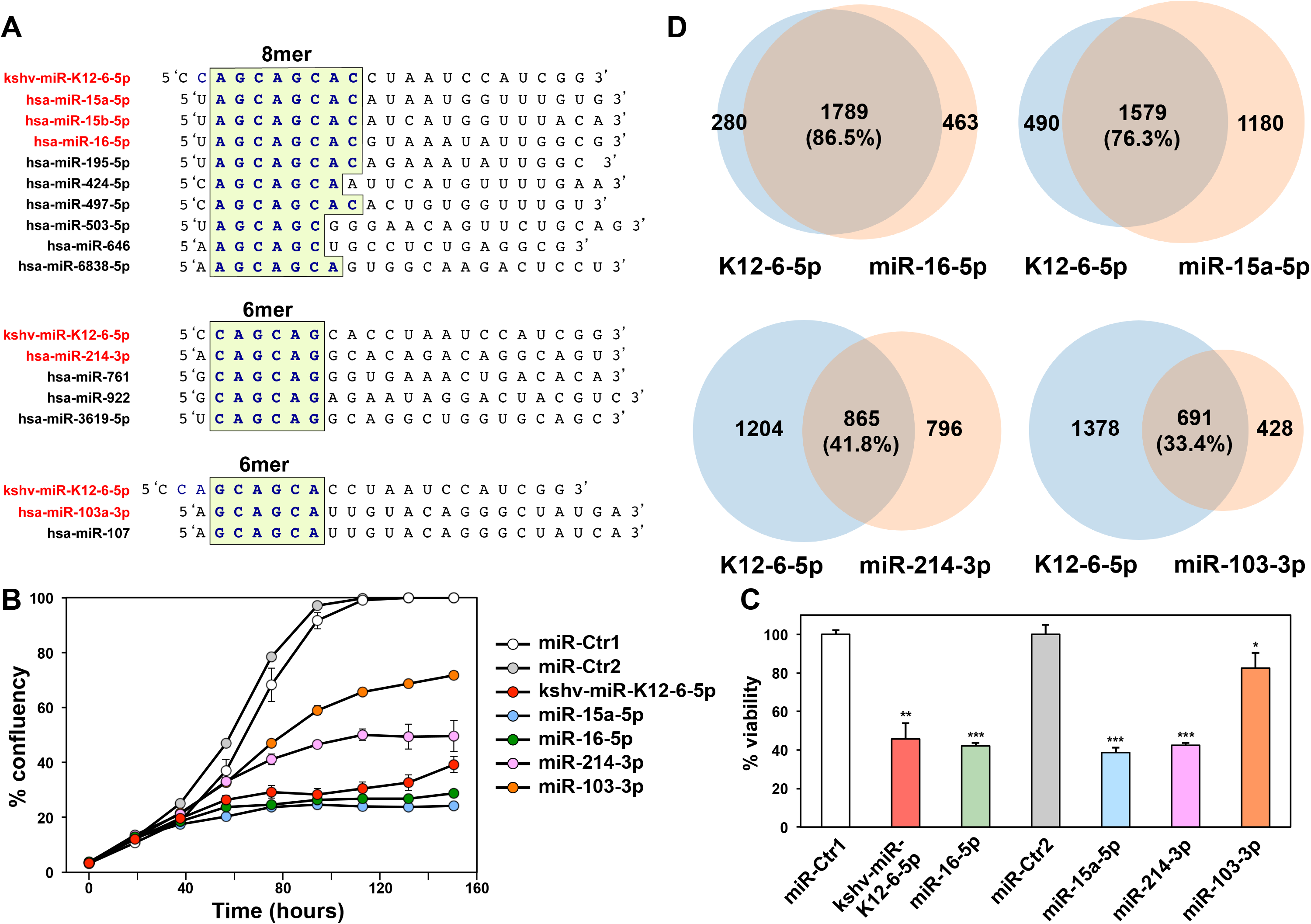
miR-K12-6-5p is most similar to the miR-15/16 family in its effects on cell survival. (A) Alignment of miR-K12-6-5p with three families of tumor suppressive miRNAs it shares seed sequences with. Identical seed sequences are highlighted. (B) Percent cell confluence over time of HeyA8 cells transfected with 10 nM of the indicated miRNAs or two miRNA non-targeting controls. Each data point represents mean ± SE of three replicates. (C) Change in viability of HeyA8 cells 96 hours after transfection with the indicated miRNAs. *** p<0.0001, ** p<0.001, * p<0.05 determined by Student’s T-test each compared to the respective control. (D) Overlap of RNAs detected by RNA-Seq downregulated in HeyA8 cells (>1.5-fold) 48 h after transfection with kshv-miR-K12-6-5p and cells transfected with either miR-15a-5p, miR-16-5p, miR-214-3p, or mR-103-3p, each compared to non-targeting miR-Ctr.

To determine whether miR-K12-6-5p could exert toxicity in cells similar to the toxicity observed with these three miRNA families, we transfected miR-15a-5p, miR-16-5p, miR-214-3p, miR-103-3p, or miR-K12-6-5p into HeyA8 cells (Figure 2B, 2C). All miRNAs substantially reduced cell growth and reduced cell viability in these cells with both miR-15a-5p and miR-16-5p being most toxic closely followed by miR-K12-6-5p. miR-103-3p was least toxic. No toxicity of any of the miRNAs was seen when Ago2 knockout HeLa cells were treated (data not shown) confirming that the effects were dependent on the activity of the RISC. To determine which of the three miRNA families miR-K12-6-5p was most similar to in its toxic activity, we compared the significantly downregulated genes in cells treated with the various miRNAs through a RNA-Seq analysis (Figure 2D). In agreement with a recent report (Morrison et al., 2019) the highest degree of overlap of downregulated genes caused by miR-K12-6-5p expression was found with miR-16-5p (76.3%) closely followed by miR-15a-5p (76.3%). A lower number of downregulated genes were shared with miR-214-3p (41.8%). miR-103-3p showed the fewest co-downregulated genes (33.4% overlap). The extent of shared downregulated genes by each miRNA correlated with the level of its toxicity suggesting that the targeted genes could be involved in cell survival.

### miR-K12-6-5p predominantly targets essential survival genes through a noncanonical 3-8 6mer seed

Not only overexpression of miR-15a-5p/16-5p caused a substantial downregulation of essential survival genes (see Figure 1F) but also miR-K12-6-5p, miR-103-3p and miR-214-3p resulted in downregulation of such genes (Figure 3A). Interestingly, with both miR-16-5p and miR-K12-6-5p we also detected a significant inverse enrichment of nonsurvival genes in the downregulated genes (Figures 1F and **3A**) confirming that these two miRNAs were most similar in their activity.

**Figure 3.**
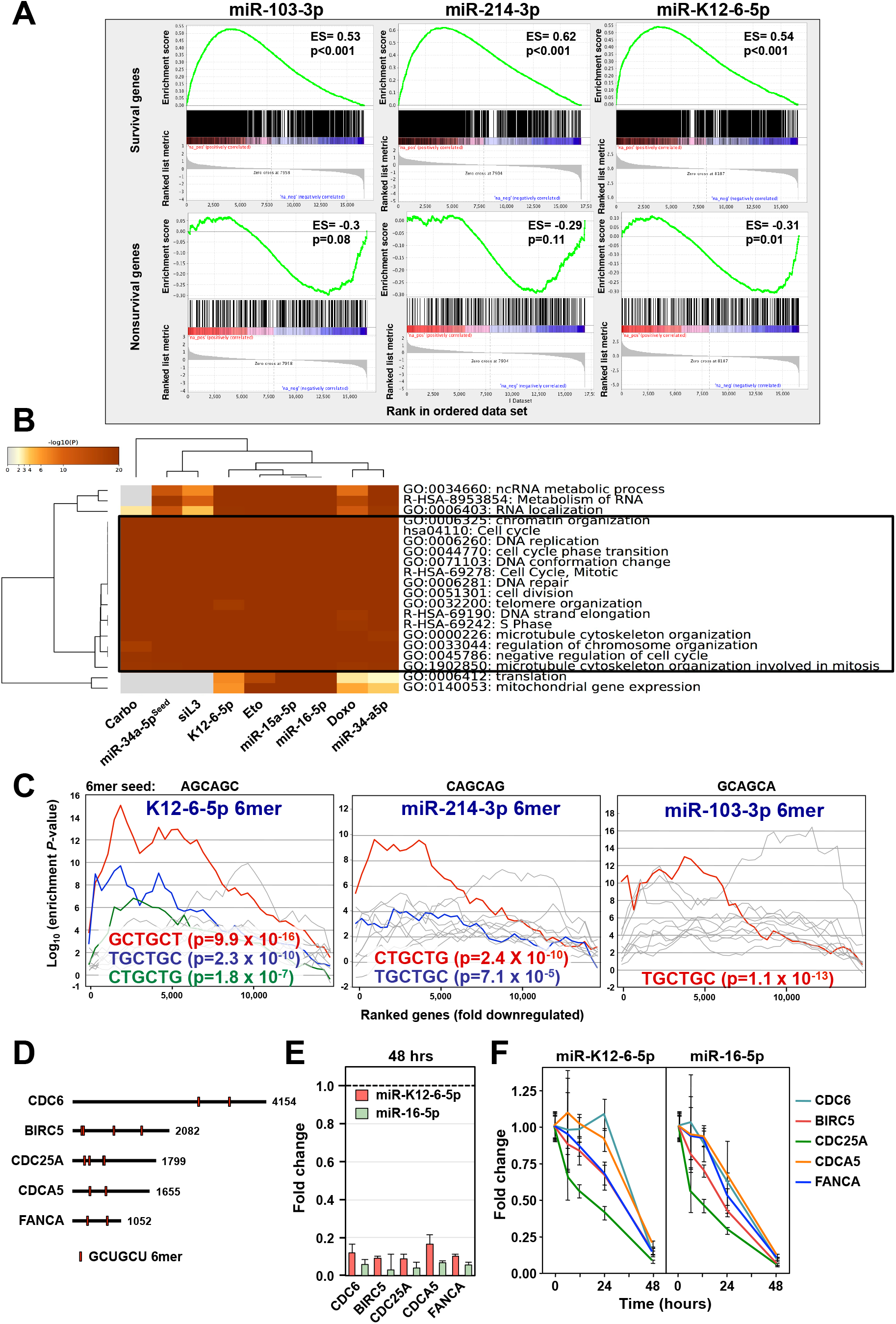
miR-K12-6-5p kills cells in part through DISE similar to miR-15/16-p. (A) Gene set enrichment analysis for a group of 1846 survival genes (top three panels) and 416 non-survival genes (bottom three panels) after transfecting HeyA8 cells with either miR-15a-5p, miR-16-5p, or si15a/16-5p^6Seed^. mod-siNT1 and a non-targeting miR-Ctr served as controls, respectively. p-values indicate the significance of enrichment and the enrichment score (ES) is given. (B) Metascape gene ontology analysis comparing the downregulated genes in cells treated with either miR-K12-6-5p, miR-15a-5p, miR-16-5p, siL3 (see (Putzbach et al., 2017)), siR-34a-5p^6Seed^, miR34a-5p, or the three genotoxic chemotherapeutic drugs (see (Gao et al., 2018)). GO terms shared by all treatments are boxed. (C) Sylamer plots for the list of either ORF or 3′UTRs of mRNAs in cells treated with either K12-6-5p (6mer seed: AGCAGC), miR-214-3p (6mer seed: CAGCAG), or miR-103-3p (6mer seed: GCAGCA) sorted from downregulated to upregulated. For each plot the ten most highly enriched seed matches are shown. The three most highly enriched ones are labeled in red, blue and green, respectively. Bonferroni-adjusted p-values are given. (D) Location of the putative GCUGCU 6mer seed matches in the 3’UTRs of the 5 tested genes (shown at scale). (E) Real time PCR quantification of the 5 genes in D in the samples of cells transfected with either miR-K12-6-5p or miR-16-5p as submitted for RNA Seq analysis. Expression levels are normalized to Crl1 transfected cells (stippled line). Data are shown with variance between duplicate biological replicates. (F) Real time PCR quantification kinetics of the same 5 genes in RNA samples isolated from HeyA8 cells transfected with either 10 nM miR-K12-6-5p or miR-16-5p. Expression levels at each time are normalized to Crl1 transfected cells. Samples are shown with StDev of three biological replicates.

To determine to what extent miR-K12-6-5p affects cell growth through 6mer seed toxicity, we performed a Metascape analysis comparing the gene ontologies of the enriched gene sets downregulated by miR-K12-6-5p and a number of stimuli we recently reported to kill cells in part through 6mer seed toxicity: a CD95L-derived toxic siRNA (Putzbach et al., 2017), miR-34a-5p (Gao et al., 2018), miR-34a-5p^6seed^ (Gao et al., 2018), and the three genotoxic chemotherapeutic reagents doxorubicin, etoposide and carboplatin (Gao et al., 2018) (Figure 3B). The genes that were downregulated in response to all these treatments had widely overlapping gene ontologies that were consistent with cells dying of 6mer seed toxicity (Gao et al., 2018; Putzbach et al., 2017; Putzbach et al., 2018b). These data suggest that miR-K12-6-5p engages a cell death mechanism that is similar to the ones triggered by the other DISE inducing reagents.

To determine which seed matches in the 3’UTRs are potentially being targeted by the v-miRNA, compared to cellular miRNAs, we performed another Sylamer analysis (Figure 3C). The most highly and most significantly enriched seed matches found in the 3’UTR of the genes downregulated in cells treated with miR-K12-6-5p was GCTGCT, the canonical 6mer seed match of miR-15a-5p/16-5p (see Figure 1D). However, this suggests that miR-K12-6-5p uses a noncanonical 6mer seed starting at position three. The most highly enriched seed matches in the 3’UTR of downregulated genes in cells transfected with either miR-214-3p or miR-103-3p also corresponded to their canonical and predicted 6mer seed starting at position two of the miRNA. Notably, there was no enrichment for miRNA targets in the open reading frames (ORFs) of the same mRNAs analyzed in Figure 3C (data not shown). These results suggest that while the four tested cellular miRNAs target mRNAs using a canonical 6mer seed, the viral miR-K12-6-5p seems to utilize a noncanonical seed located at position 3-8 consistent with a recent study (Morrison et al., 2019).

As shown in Figure 2B cells started to slow down in growth around 40 hours after transfection with either miR-K12-6-5p or miR-16-5p. To determine whether the targeting of a set of essential survival genes by these miRNAs could be causing this, we identified the 5 most highly expressed genes that contain at least two putative GCUGCU seed matches in their 3’UTR and that were downregulated in cells transfected either with K12-6-5p or with miR-16-5p. Three of these genes are established cell cycle regulators (CDC6, CDC25A, CDCA5), one is involved in DNA repair (FANCA), and one codes for an antiapoptotic protein (BIRC5) (Figure 3D). First, we validated by qPCR that they were downregulated in the RNA subjected to the RNA Seq analysis (Figure 3E). Then we performed kinetics starting with very early time points of 6 and 12 hrs to determine how long before any detectable cell death these mRNAs were downregulated. The majority of the genes showed clear signs of silencing very early after miRNA transfection and they were again all efficiently silenced at 48 hrs (Figure 3F), a time point at which no dying cells could be seen (**Movies S1-S3**). This analysis suggests that the knockdown of these genes is the cause of the death of the cells rather than the result of dying cells.

### Many v-miRNAs can regulate cell fate through 6mer seed toxicity using a noncanonical 6mer seed

Our data suggest that the v-miRNA miR-K12-6-5p negatively regulates cell growth at least in part through 6mer seed toxicity similar to the endogenous miRNAs of the miR-15/16-5p family. We had identified the most toxic 6mer seeds by screening the two human cancer cell lines HeyA8 (ovarian cancer) and H460 (lung cancer) (6merdb.org) (Gao et al., 2018). To test whether other v-miRNAs would be cytotoxic to cells through 6mer seed toxicity and to test whether they also used noncanonical seeds, we transfected 215 mature v-miRNAs coded for by 17 human pathogenic viruses listed in miRBase at a concentration of 25 nM into the two cell lines and quantified cell death using a cell viability assay (**Table S2**). We chose the same two cell lines we had used in our previous screen to directly compare the results and also to isolate the tissue/host cell independent activities of each miRNA. We ranked the 215 v-miRNAs according to the average toxicity in the two cell lines (Figure 4A). The congruence between the results of the two cell lines was as high as the one we reported for the toxicity of the 4096 seed containing siRNAs (**Figure S1A**; r=0.64 and 0.68, respectively) suggesting that the v-miRNAs were toxic through a mechanism independent of host cell specific viral functions.

**Figure 4.**
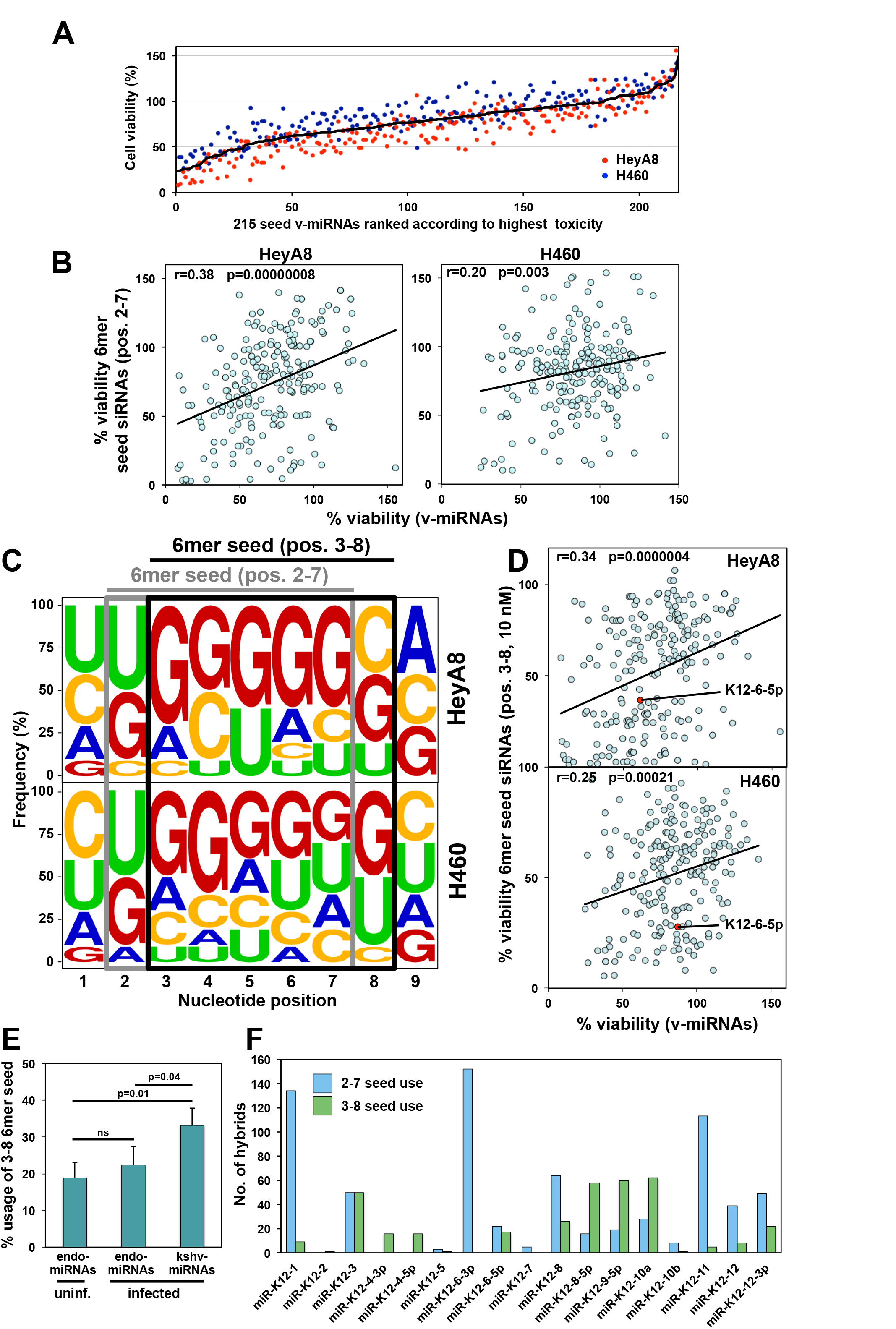
Evidence for 6mer seed toxicity across 215 v-miRNAs encoded by human pathogenic viruses. (A) Results of the 215 v-miRNA screen in two human cell lines. Cells were reverse transfected in triplicates in 384 well plates with 25 nM of individual v-miRNAs. The cell viability of each 6mer seed duplex was determined by quantifying cellular ATP content 96 hours after transfection. The v-miRNAs are ranked by the average effect on cell viability of the two cell lines from the most toxic (left) to the least toxic (right). (B) Regression analysis showing correlation between the toxicity observed in the human lung cancer cell line H460 (left) and the human ovarian cancer cell line HeyA8 (right) 96 hours after transfection with either 25 nM 215 v-miRNAs or 25 nM 6mer seed siRNAs carrying the same 6mer seed in a neutral backbone. p-Values were calculated using Pearson correlation analysis. (C) Nucleotide composition at each of the first 9 nucleotide positions in the top 10 most toxic v-miRNAs transfected into HeyA8 (top) or H460 (bottom) cells. The canonical 6mer seed (position 2-7) and the noncanonical 6mer seed (position 3-8) are boxed in grey and black, respectively. (D) Regression analysis showing correlation between the toxicity observed in H460 (top) and HeyA8 (bottom) cells 96 hours after transfection with either 25 nM 215 v-miRNAs or 10 nM of the siRNAs carrying matching 6mer seeds (position 3-8) tested previously (Gao et al., 2018). p-values were calculated using Pearson correlation analysis. (E) Percent use of the noncanonical 6mer seed (position 3-8) by either cellular miRNAs or v-miRNAs that were identified to be part of CLASH hybrids in an analysis of endothelial cells uninfected or infected with KHSV (Gay et al., 2018). StDev is given based on triplicate data sets. Significance was calculated using two-sample Student’s t-test. ns, not significant. (F) All hybrids detected in the three triplicates of the CLASH analysis of KHSV infected cells that contain a kshv-miRNA with full complementarity between either its canonical 6mer seed (pos. 2-7) or its noncanonical seed with a piece of mRNA (pos. 3-8).

To assess the contribution of just the 6mer seed to the activity of each v-miRNA, we performed a screen with siRNAs each carrying one of the 196 different 6mer seeds present in the 215 v-miRNAs in a neutral backbone (**Table S3**). Cells were transfected with 25 nM siRNA. The data on the 196 siRNAs of this screen correlated well, especially in the HeyA8 cells, with the 196 siRNAs that were part of the previous screen of the 4096 6mer seed siRNAs transfected at 10 nM (Gao et al., 2018) (**Figure S1B, Table S4**). Most importantly, regression analysis showed a strong correlation in both cell lines between the toxicity observed with either 25 nM 215 v-miRNAs or 25 nM 6mer seed siRNAs carrying the same 6mer seed in a neutral backbone (Figure 4B), suggesting that the first 6 nucleotides of the seed present in the v-miRNAs were important to the exertion of toxicity in the cells.

The most toxic 6mer seeds are rich in Gs with a preferential enrichment for Gs at the 5’ end of the 6mer seed (Gao et al., 2018). To determine the nucleotide composition of the most toxic seeds in the v-miRNAs, we plotted the relative abundance of the four nucleotides in each of the first 9 positions (the extended seed region) of the ten most toxic v-miRNAs (~5% of all tested v-miRNAs) in the screen in both cell lines (Figure 4C). While v-miRNAs in general were found to have a slightly elevated G content in the 6mer seed region (**Figure S2A**), similar to the 4096 6mer seed screen, the most toxic v-miRNAs were strongly enriched in Gs (Figure 4C, grey box). Curiously however, G was not the most abundant nucleotide in the first position of this 6mer sequence. Based on our analysis and the recent report on miR-K12-6-5p and its potential of using a noncanonical 6mer seed starting at position three (Morrison et al., 2019), we realized that the optimal seed composition for the most toxic 6mer seeds we had reported could be achieved by shifting the seed by one nucleotide to start at position three (black box in Figure 4C). In fact, for the HeyA8 cells this resulted in the generation of a seed composition remarkably similar to the one we had reported for the 200 (~5%) most toxic seeds in the 4096 seed siRNA screen (**Figure S2B**) (Gao et al., 2018). Our data suggest, firstly, that many v-miRNAs indeed have the capacity to affect cell fate through 6mer seed toxicity and secondly, and similar to miR-K12-6-5p, that many v-miRNAs may use a noncanonical toxic 6mer seed that starts at position three of the mature miRNA. To test this assumption, we plotted the toxicity of the 215 v-miRNAs versus that of the 6mer seed siRNAs tested in our previous screen that matched positions 3-8 in the v-miRNAs (representing 197 different 6mer seeds, **Table S5**) (Figure 4D). The correlation was stronger and more significant than when compared with the original screen results of the seed containing siRNAs that matched the 196 canonical seeds (**Figure S1C**). The data suggest that many v-miRNAs can kill cells through use of a noncanonical 6mer seed.

To explore to what degree the canonical versus a noncanonical 6mer seed sequence contributes to the toxicity of v-miRNAs across both cell lines, we compared the screen data of two groups of v-miRNAs for correlations between sequence identity and functional toxicity in cells. Among the 215 tested v-miRNAs we found 6 pairs of v-miRNAs that are identical in sequence, 13 pairs of v-miRNAs (including one triplet) that share the same 2-7 6mer seed, and 10 pairs (including two triplets) that share the same 3-8 6mer seed, both with diverging sequences outside the seed sequence (**Table S3**). When plotting every pair and triplet of v-miRNAs against each other as expected, the v-miRNAs with complete sequence identity affected cell viability in a very similar way (**Figure S1D**, top left panel). When plotting the v-miRNA pairs that only shared a canonical 6mer seed, we still found a significant correlation in viability (r=0.48 in **Figure S1D**, top right panel). However, this correlation was more pronounced and more significant when the v-miRNA pairs were analyzed that shared a noncanonical 6mer seed (r=0.56 in **Figure S1D**, bottom left panel).

Many studies have shown a compensatory role of nucleotides beyond the canonical seed through supplementary pairing (Broughton et al., 2016; McGeary et al., 2019). One such supplementary pairing region involves position 13-16 of the miRNA (Figure 5A) (Sheu-Gruttadauria et al., 2019). An involvement of supplementary seed pairing in the toxicity of K12-6-5p was unlikely as the sequence of K12-6-5p shared no identity with the modified backbone carrying the AGCAGC 6mer seed that corresponded to the 3-8 seed in K12-6-5p (bottom two sequences in Figure 5A). This modified siRNA was as toxic as K12-6-5p itself (compare Figure 1B and Figure 2C). However, to address a possible contribution of this supplementary pairing region directly, we separated the data of all v-miRNAs in the screen into a group that while sharing an identical 6mer seed either shared the same sequence or differed in the supplementary region. Interestingly, the correlation of the miRNAs that carry the same 6mer seed but differ in position 13-18 was greater and more significant than the correlation of the miRNA pairs that share position 13-16 (**Figure S1D**, bottom right panel). This supports our conclusion that it is predominantly the 6mer seed that determines toxicity with little contribution of supplementary seed pairing to this phenomenon. These analyses confirm that while the entire length of the v-miRNAs can have an effect on cell growth, a substantial part of this toxicity is dictated by only the 6 nucleotides in the 6mer seed and this is more pronounced when the noncanonical 6mer seed is considered.

**Figure 5:**
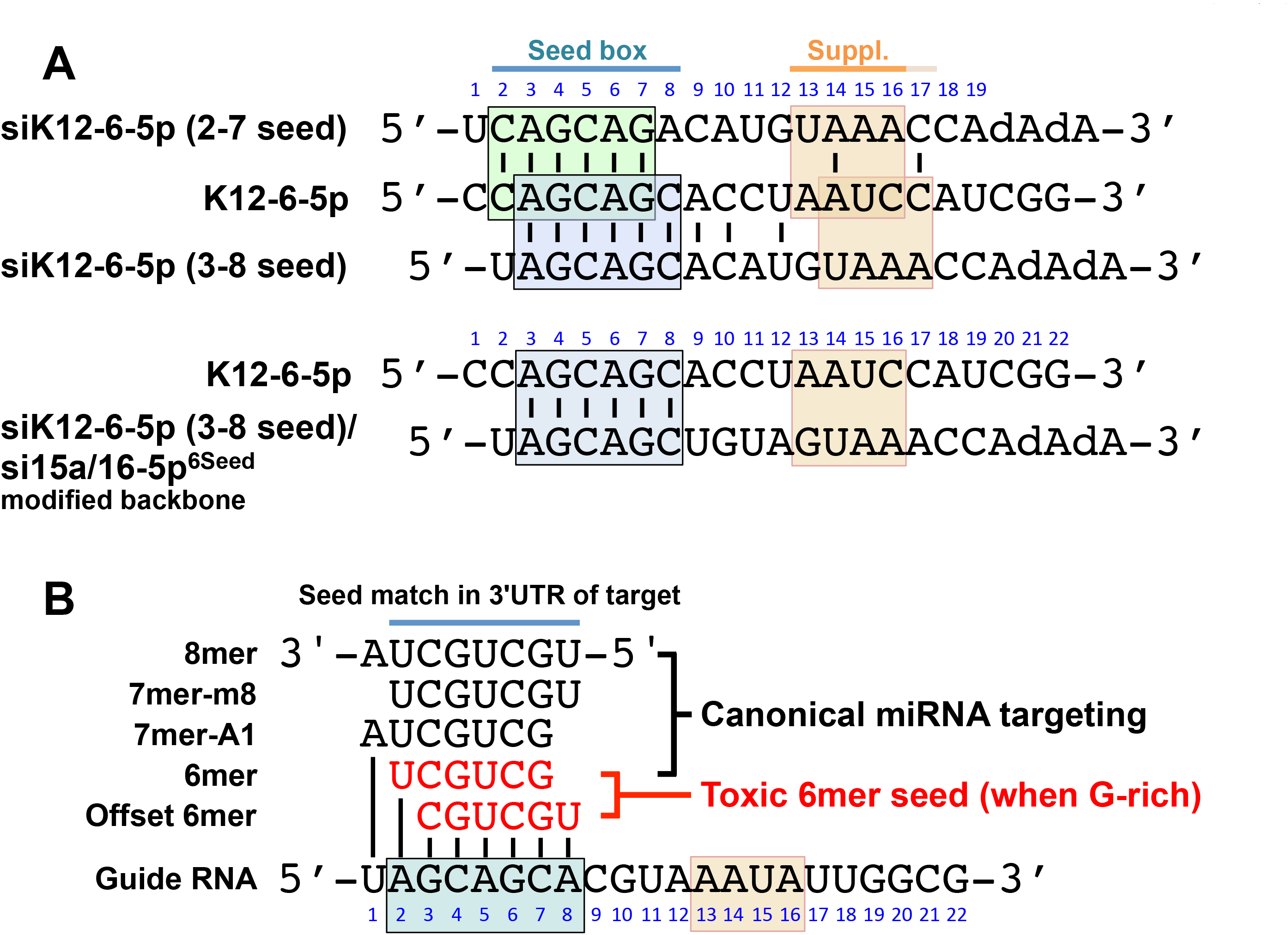
Canonical and noncanonical 6mer seed usage in the context of miRNA seeds. (A) Alignment of K12-6-5p (second line) with the canonical (top line) or the noncanonical (third line) K12-6-5p seed embedded in the original backbone of siRNAs used in the screen of 4096 6mer seeds (Gao et al., 2018). Below this alignment K12-6-5p is shown with its noncanonical seed aligned with the modified backbone shown in Figure 1A. Note, between these two sequences there is no identity outside the 6mer seed box including in the supplementary pairing region (suppl.). (B) Location of the 6mer seeds relative to the longer seed boxes usually found in miRNAs (shown is miR-16-5p). The 6mer seed sequences can be part of longer seed boxes. Longer seeds allow more selective and more profound targeting but an embedded 6mer seed when G rich (either canonical or offset) can target essential survival genes with low efficiency but to a degree enough to be toxic to cells. This figure was modified after (Riffo-Campos et al., 2016).

To determine whether the 3-8 noncanonical toxic 6mer seed was predominantly used by miRNAs of specific viruses, we separated the data into individual viruses and correlated the toxicity associated with canonical and non-canonical seed of the v-miRNA with activity of the canonical seed in the corresponding siRNA (**Figures S3** and **S4**). We only found two groups of viruses that showed a significant change in correlation in both cell lines between the 2-7 and the 3-8 toxic 6mer seed use. A combined group of viruses that each code for two or less miRNAs (BKV, BPCV1, BPCV2, JCV, SV40 and TTV) seems to prefer the canonical 2-7 seed (green boxes in **Figures S3** and **S4**). In contrast, the miRNAs encoded by KSHV showed a significantly increased correlation in HeyA8 cells and a correlation almost reaching significance in H460 cells when plotting the noncanonical 3-8 seed (red boxes in **Figures S3** and **S4**).

Finally, to validate our results with data on cells infected with a relevant virus that carries multiple miRNAs, and to break down the use of the 3-8 seed in the KSHV miRNAs, we analyzed a data set on KSHV infected endothelial cells generated by using a modified Cross-linking, Ligation, and Sequencing of Hybrids (CLASH) method (Gay et al., 2018). CLASH allows for the unambiguous identification of miRNAs and their targets by ligating the two species while bound to the RISC. We extracted the CLASH hybrid data from two triplicate samples that had been generated using either uninfected cells or cells infected with KSHV. Only two types of hybrids were counted: the ones that showed complete complementarity between the seed match in the target and positions 2-7 of the miRNA (the canonical 6mer seed) and the ones that had this region of complementarity shifted to position 3-8 (the noncanonical 6mer seed). In each case the nucleotides preceding the seed and the nucleotide following the seed had to be mismatched to focus the analysis on the two types of 6mer seeds. We then plotted the occurrence of a noncanonical 6mer seed in percent of total (Figure 4E). When analyzing the cellular miRNAs in the uninfected cells, about 20% of the detected seed interactions were noncanonical (Figure 4E, first column). A similar proportion was found in the cellular miRNAs in the virus infected cells (Figure 4E, second column). However, the hybrids involving v-miRNAs in the virus infected cells showed a significantly higher use of the noncanonical 3-8 6mer seed (Figure 4E, third column). Of the 25 mature miRNAs coded for by KSHV, 17 were detected in CLASH hybrids. To determine the 6mer seed use of individual KHSV miRNAs, we analyzed all hybrids obtained from virus infected cells (Figure 4F). While K12-6-3p showed no evidence of noncanonical 6mer seed use, other miRNAs including miR-K12-3 and miR-K12-6-5p showed a substantial use of the 3-8 6mer seed. Three miRNAs (miR-K12-8-5p, miR-K12-9-5p and miR-K12-10a) were found in hybrids that predominantly made use of the 3-8 6mer seed. In summary, we provide first evidence that v-miRNAs can engage the 6mer seed toxicity mechanism and that many of them do this through a noncanonical use of the 6mer seed.

## DISCUSSION

We recently identified RNAi-based 6mer seed toxicity as an ancient kill code embedded in the genome (Gao et al., 2018). It can kill cells independent of tissue of origin and species. This mechanism is based on only a 6mer seed sequence and the most toxic seeds were found to be G-rich targeting C-rich seed matches in the 3’UTR of essential survival genes. We also provided evidence that this mechanism may be at least 800 million years old. Based on these findings we predicted that viruses coevolved with this fundamental mechanism and either avoid G-rich seeds in small RNAs such as miRNAs or use them to kill cells.

After reporting data to suggest that about 80% of the toxic activity of the master tumor suppressive miRNA miR-34a-5p is based on the first 6 nucleotide of its seed (Gao et al., 2018), we now tested another major tumor suppressive miRNA family, miR-15/16-5p, and found 6mer seed toxicity also significantly contributes to its toxic activity. While one of the most prominent targets for miR-15/16 is the antiapoptotic protein Bcl-2 (Cimmino et al., 2005), many of its targets are cell cycle regulators (Calin et al., 2008; Linsley et al., 2007; Liu et al., 2008) and a recent analysis showed that miR-16 downregulated genes are generally involved in proliferation (Morrison et al., 2019). This type of targeting is consistent with miR-15/16 affecting cell growth and viability through 6mer seed toxicity. Consistently, we found that >55% of the toxic activity of miR-15/16-5p comes from its 6mer seed. This is quite high considering that a substantial number of targets of miR15b/16 are actually targeted through complementary between the 3’UTR and the 3’ end of the miRNA (Helwak et al., 2013).

Because it was recently shown that the v-miRNA kshv-miR-K12-6-5p is an oncomimetic of miR-15/16-5p, we were wondering whether v-miRNAs such as miR-K12-6-5p could use the 6mer seed kill code to affect cell fate. Our analysis of the activity of miR-K12-6-5p compared to its related cellular miRNAs miR-15/16-5p, miR-214-3p, and miR-103a-3p/107, all of which share sequence identities in their seed sequences with miR-K12-6-5p, confirmed and extended the recent finding that miR-K12-6-5p functionally is most similar to miR-15/16-5p (Morrison et al., 2019) by demonstrating that much of the toxicity of both miR-15/16-5p and miR-K12-6-5p relies on only 6 nucleotides in their seed sequences. Indeed, Morrison et al. (2019) reported in the context of KSHV-infected tumor cells that miR-K6-5p targets cell cycle regulators and genes involved in cell proliferation consistent with our data that this miRNA negatively regulates proliferation or survival through 6mer seed toxicity.

When testing all 4096 possible 6mer seed combinations in our previous screen, for practical reasons we had to design an artificial backbone of an siRNA duplex in which to test all seeds. None of these siRNAs therefore represented actual miRNAs found in cells. Another caveat was that the "neutral" backbone we had designed may have obscured the results by biasing the targeting of certain genes that carry sequences complementary to backbone regions. The new screen with 215 viral miRNAs now allowed us to test the 6mer seed toxicity concept using a set of miRNAs carrying different 6mer seeds all in their physiologically relevant and mostly different backbones. Considering that v-miRNAs in particular were shown before to not only target by using canonical 5’ seed sequences, but also by using 3’ seeds, or sequences located in the center of the miRNA (Gay et al., 2018), our findings of a substantial correlation (20-38%) between the toxicity of the tested v-miRNAs and the toxicity of the matching siRNAs just containing the same 6mer seed sequence points at 6mer seed toxicity as a robust cellular mechanism to regulate cell fate by miRNAs. This level of correlation is remarkable when one considers that not all v-miRNAs (or any miRNA) will be toxic to cells through only 6mer seed toxicity. Many miRNAs will likely be toxic through conventional seed targeting of individual genes involving seeds longer than 6 nucleotides and sequences outside the 5’ seed box.

By studying the miR-15/16-5p mimic miR-K12-6-5p we and others (Morrison et al., 2019) noted that while miR-K12-6-5p is functionally most similar to the miR-15/16-5p family, the seed most likely used by miR-K12-6-5p is shifted by one nucleotide starting at position 3. A noncanonical seed usage beginning at position 3 was first described for the miRNA bantam in Drosophila (Nahvi et al., 2009). Interestingly, by testing the toxicity and 6mer seed composition of the 215 v-miRNAs we now provide evidence to suggest that the use of such a noncanonical 6mer seed by v-miRNAs is more frequent than by cellular miRNAs. A substantial use of a noncanonical 6mer seed was seen for miR-K3. miR-K3 shares a perfect offset 7mer seed with miR-23 (Manzano et al., 2013). Despite the offset seed, both miR-K3 and miR-23 target similar genes just as K12-6-5p shares many targets with miR-15/16-5p. While miR-K3 has a few unique targets which are not shared by miR-23 (Grosswendt et al., 2014; Manzano et al., 2013), it was recently reported that the noncanonical seed may not be functionally relevant (Grosswendt et al., 2014). The authors selected five noncanonical miR-K3 interactions for further examination of their regulatory potential in reporter assays but did not detect reporter repression mediated by any of these sites (Grosswendt et al., 2014). Our data now suggest that the noncanonical 6mer seed may be regulating cell survival in many of these v-miRNAs through DISE. This type of targeting is different from canonical miRNA targeting that relies on seed sequences in the miRNAs that are longer than 6 nucleotides and additional sequences outside the seed (Figure 5B) (Broughton et al., 2016). The more nucleotides are involved in pairing with the mRNA target the more selective and more profound the downregulation will be. In contrast, targeting by a short seed of only 6 nucleotides decreases selectivity of targeting and likely results in the downregulation of a larger number of genes each to a lower extent. It is likely, that targeting hundreds of essential survival genes simultaneously when the 6mer seed is G-rich (each of which is defined to be an essential gene individually (Putzbach et al., 2017)) at a relative low level should be highly toxic to cells.

Remarkably, the nucleotide composition of the noncanonical 6mer seed of the 5% most toxic v-miRNAs was almost identical to the 5% most toxic canonical seeds identified in the 4096 6mer seed screen. In addition, the toxicity of the tested v-miRNAs when considering the noncanonical 3-8 6mer seed correlated much better with the results of the 4096 siRNA screen than when considering the canonical 2-7 6mer seed. Our data suggest that during evolution certain v-miRNAs evolved to use 6mer seed toxicity to regulate cell growth, cell cycle and survival of their host cells. It has been proposed that seed shifting is a mechanism of miRNA adaptation during evolution to allow acquisition of divergent function (Berezikov, 2011; Bofill-De Ros et al., 2019). The small but significant increased use of a noncanonical seed by v-miRNAs may be a reflection of an ongoing evolutionary process that was conserved in the v-miRNAs at the time the miRNAs were acquired from the host cells. Future studies will be aimed at elucidating the mechanism of the alternative processing and/or seed usage of viral miRNAs and at determining whether v-miRNAs and other small virus encoded RNAs affect cell fate through 6mer seed toxicity in infected cells.

### Limitations of the Study

The viruses that code for the miRNAs tested in our screen infect different cell types and have complex life cycles. However, our results will likely be relevant to the study of the function of viral miRNAs because the rules that govern 6mer seed toxicity are tissue independent. Furthermore, the specificity of the effects of a virus on cells is often determined by the entry mechanism rather than by differential effects of the viral genes on specific host cells. An example are the studies that utilized VSVg pseudotyped HIV-1 to explore general mechanisms of viral replication and responses of CD4 negative cells to HIV-1 infection (Aiken, 1997; Akari et al., 1999; Hulme et al., 2011). To test 215 miRNAs encoded by 17 different human viruses each in a virus relevant cellular context (proper host cells and models or lytic versus latency cycle), would not have been feasible. We therefore decided to perform the screen in the same two human cancer cell lines we had used to determine the toxicity of all 4096 6mer seeds (Gao et al., 2018). As a result, the screen of viral miRNAs also allowed us to further validate the 6mer seed concept.

## AUTHOR CONTRIBUTIONS

A.E.M. and M.E.P. planned the experiments. A.E.M. and S.C., J.V. performed experiments. E.T.B. and M.J.S. performed data analyses, S.G. contributed expertise on viral miRNAs, and M.E.P. directed the study and wrote the manuscript.

## ACKNOWLEDGEMENT

We are grateful to Dr. Eva Gottwein for sharing data on miR-K6-5p prior to publication and for critically reading the manuscript. We would like to thank Ashley Haluck-Kangas for editing the manuscript. This work was funded by grant R35CA197450 to M.E.P.

