## Supplementary material for "6mer seed toxicity in viral microRNAs": Methods and supplemental figures

### Transparent Methods

#### Reagents, cell lines and antibodies

HeyA8 (RRID:CVCL_8878) and H460 (ATCC HTB-177) cells were cultured in RPMI1640 medium (Cellgro Cat#10-040) supplemented with 10% FBS (Sigma Cat#14009C), 1% penicillin/streptomycin (Mediatech Inc.), and 1% L-Glutamine (Corning Cat#25-005). Cell lines were authenticated by STR profiling.

#### siRNA and v-miRNA screens

Analysis of v-miRNAs deposited at miRbase 22.1 identified 216 different v-miRNAs encoded by 17 different viruses that can be pathogenic to humans (**Table S1**). 215 of them were synthesized into one 384 plate (MIRNA MIMIC 2.0, LIB, 0.25nmol, ThermoFisher). The library of 4096 6mer seed siRNAs were previously described (Gao et al., 2018). 196 siRNAs containing the 6mer seeds that are part of the 215 v-miRNAs were retested on HeyA8 and H460 cells. All RNA duplexes were screened in 384 well plates as described with one modification: they were reverse transfected at a final concentration of 25 nM. All data were normalized to lipid only on each plate.

#### Transfection with short oligonucleotides

50 μl transfection mix with 0.15 μl RNAiMAX and 10 nM siRNAs or miRNAs were plated. Then cells were added in 200 μl antibiotic free medium in a 96 well plate at 1000 cells/well. Custom siRNA oligonucleotides were ordered from integrated DNA technologies (IDT) and annealed according to the manufacturer's instructions. In addition to the 4096 siRNAs of the screen the following siRNA sequences were used:
siNT1modified sense (siCtr): mUmGrGrUrUrUrArCrUrArCrArCrGrArCrUrArUTT;
siNT1modified antisense (siCtr) : rArUrArGrUrCrGrUrGrUrArGrUrArArArCrCrAAA;
miR-15a/16-5p$^{6Seed}$ sense: mUmGrGrUrUrUrArCrUrArCrArGrCrUrGrCrUrUTT;
miR-15a/16-5p$^{6Seed}$ antisense: rArArGrCrArGrCrUrGrUrArGrUrArArArCrCrAAA;
The following miRNAs and negative controls were used: hsa-miR-15a-5p (Ambion, Cat. No# AM17100, Part number PM10235), hsa-miR-16-5p (mirVana® miRNA mimic, Ambion, Cat. No# AM17100, Part Number MC10339); hsa-miR-214-3p (Ambion, Cat. No# AM17100, Part Number PM12124); miR-103-3p (Ambion, Cat. No# AM17100, Part Number PM10632); kshv-miR-K12-6-5p (Ambion, Cat. No# 4464066; Assay ID MC10307), and negative controls and miR-control#1 (mirVana™ miRNA Mimic, Negative Control #1, Ambion Cat. No# 4464058), and miR-control#2 (miRNA precursor negative controls #1, Ambion, Cat. No# AM17110).

#### Monitoring cell growth by IncuCyte and cell death assays

Cells were seeded in a 96-well plate in triplicates. The plate was then scanned using the IncuCyte ZOOM live-cell imaging system (Essen BioScience). Images were captured every four hours using a 10× objective. Cell confluence was calculated using the IncuCyte ZOOM software (version 2015A). To assess cell viability, treated cells were subjected to a quantification of nuclear fragmentation or ATP content. To measure the cellular ATP content, cells were reverse transfected with siRNAs in a 96 well plate at 1000 cells per well. 96 hours after transfection, media in each well was replaced with 70 μl of fresh media and 70 μl of CellTiter-Glo cell viability reagent (Promega). The plates were shaken for 5 min and incubated at room temperature for 15 min. Luminescence was then read on the BioTek Cytation 5.

#### RNA-Seq analysis

For RNA-Seq data 50,000 cells/well HeyA8 cells were reverse transfected in duplicate in 6-well plates with 10 nM of siRNAs or miRNAs. Cells were lysed 48 hours after transfection using Qiazol. Total RNA was isolated using the miRNeasy Mini Kit (Qiagen, Cat.No# 74004) following the manufacturer's instructions. An on-column digestion step using the RNAse-free DNAse Set (Qiagen, Cat.No# 79254) was included for all RNA-Seq samples. RNA libraries were generated and sequenced (Genomics Core Facility at the University of Chicago). The quality and quantity of the RNA samples were checked using an Agilent bio-analyzer. Paired end RNA-SEQ libraries were generated using Illumina TruSEQ TotalRNA kits using the Illumina provided protocol (including a RiboZero rRNA removal step). The libraries were sequenced on an Illumina HiSEQ4000 using Illumina provided reagents and protocols. Adaptor sequences were removed from sequenced reads using TrimGalore (https://www.bioinformatics.babraham.ac.uk/projects/trim_galore), The trimmed reads were aligned to the hg38 version of the human genome, using STAR v2.5.2. Aligned reads were associated with genes using HTSeq v0.6.1, and the UCSC hg38 transcriptome annotation from iGenomes. Differentially expressed genes were identified using the edgeR R package.

**Quantitative real-time PCR.** Total RNA extraction, RNA quantification and cDNA preparation were done as previously described (Qadir et al., 2017). 200 ng of RNA in 20 μl of cDNA reaction volume was used to generate cDNA using the High-Capacity cDNA Reverse Transcription Kit (Applied Biosystems). The expression of CDC25A (Hs00947994_m1), BIRC5 (Hs04194392_s1), CDC6 (Hs00154374_m1), FANCA (Hs01116668_m1), and CDCA5 (Hs00969392_g1) was quantified using specific primers from Life technologies. The expression levels of target genes were normalized to the reference housekeeping gene, GAPDH (Hs00266705_g1). Fold differences were then calculated for each treatment group using normalized $C_T$ values for the control. The qPCR was run in 96 well qPCR plates with 20 µl of reaction mixtures in AB 7500 Fast Real-Time system in QuantStudio Real-Time PCR System from Applied Biosystems.

**Data analyses**

GSEA was performed using the GSEA software version 3.0 from the Broad Institute downloaded from https://software.broadinstitute.org/gsea/. A ranked list was generated by sorting genes according the $Log_{10}$(fold downregulation). The Pre-ranked function was used to perform GSEA using the ranked list. 1000 permutations were used. Default settings were used. The ~1800 survival genes and ~420 non-survival genes defined previously (Putzbach et al., 2017) were used as custom gene sets. Default settings were used.

Sylamer analysis (van Dongen et al., 2008) was used to find enrichment of small word motifs in the 3'UTRs of genes enriched in those that are most downregulated. 3'UTRs or ORFs were used from Ensembl, version 76. For each gene the longest transcript was used as described previously (Putzbach et al., 2017). As required by Sylamer, they were cleaned of low-complexity sequences and repetitive fragments using respectively Dust (Morgulis et al., 2006) with default parameters and the RSAT interface (Medina-Rivera et al., 2015) to the Vmatch program, also run with default parameters. Sylamer (version 12–342) was run with the Markov correction parameter set to four. Bonferroni adjusted p-values were calculated by multiplying the unadjusted p-values by the number of permutations for each length of word searched for.

The GO enrichment analyses shown in Figure 1D and 1E were performed using the GOrilla GO analysis tool at http://cbl-gorilla.cs.technion.ac.il using default setting using different p-value cut-offs for each analysis. GO analyses across multiple data sets (Figure 3B) were performed using the software available on www.Metascape.org and default running parameters.

Venn diagrams were generated using http://bioinformatics.psb.ugent.be/webtools/Venn/ using default settings.

Density plots showing the contribution of the four nucleotides G, C, A and U at each of the 6mer seed positions were generated using the Weblogo tool at http://weblogo.berkeley.edu/logo.cgi using the frequency plot setting.

Analysis of qCLASH data. Data sets on qCLASH analysis of endothelial cells uninfected or infected with KHSV were downloaded at GSE101978.The structures of miRNA/target interactions were predicted from Clash hybrid sequences using the program RNAfold (http://rna.tbi.univie.ac.at/cgi-bin/RNAWebSuite/RNAfold.cgi) with default parameters. The resulting Vienna diagrams were sorted into those with a mismatch in the first position followed by 6 exact matches, encoded as .((((((. and those with two mismatches followed by 6 exact matches, encoded as ..((((((. , henceforth referred to as a 2-7 or 3-8 seed match, respectively. In both cases, a trailing mismatch was required at the 3' end. To determine the proportion of noncanonical seed 6mer use, the ratio of the number of hybrids containing a canonical or a noncanonical seed was calculated.

**Statistical analyses**

Continuous data were summarized as means standard errors and dichotomous data as proportions. Continuous data were compared using t-tests for two independent groups. For evaluation of continuous outcomes over time, two-way ANOVA was performed using the Stata 14 software with one factor for the treatment conditions of primary interest and a second factor for time treated as a categorical variable to allow for non-linearity. Comparisons of single proportions to hypothesized null values were evaluated using binomial tests. Pearson correlation coefficients (r) and p - values as well as Wilcoxon rank test were calculated using StatPlus (v. 6.8.1).

**Data availability**

RNA sequencing data generated for this study is available in the GEO repository: GSE135453 (https://www.ncbi.nlm.nih.gov/geo/query/acc.cgi?&acc=GSE135453, reviewer access token: alexaukinnavbkj). All source data are available upon request.

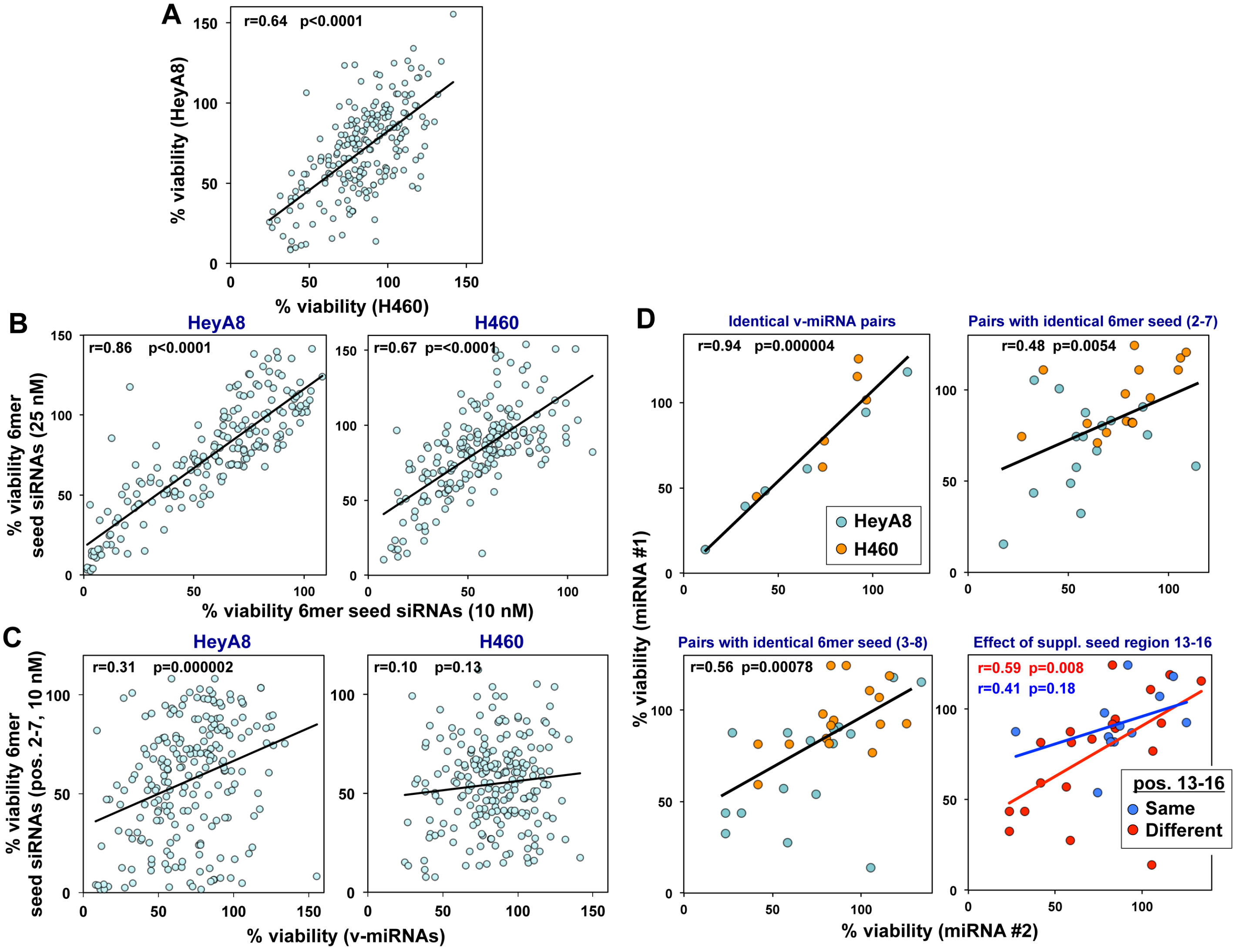


**Figure S1. Related to Figure 4. Correlation between toxicity exerted by v-miRNAs and their 6mer seeds.** (A) Regression analysis showing correlation between the toxicity observed in H460 and HeyA8 cells 96 hours after transfection with 215 v-miRNAs at 25 nM. (B) Regression analysis showing the correlation in cell viability between the 196 siRNAs that carry the same 6mer seed as the 215 human v-miRNAs of the original screen (at 10 nM, x-axis) and the new screen (at 25 nM, y-axis). (C) Regression analysis showing the correlation between the toxicity observed in HeyA8 (top) and H460 (bottom) cells 96 hours after transfection with either 25 nM of the 215 v-miRNAs or 10 nM of the siRNAs carrying matching 6mer seeds (position 2-7) tested previously (Gao et al., 2018). (D) Regression analysis showing pairwise comparison of two v-miRNAs in a group of v-miRNAs identical in sequences over the entire length (top left), a group of v-miRNAs only sharing a canonical 6mer seed (position 2-7, top right), and a group of v-miRNAs sharing a noncanonical 6mer seed (position 3-8, bottom left). Bottom right: Regression analysis showing pairwise comparison of two v-miRNAs in a group of v-miRNAs with an identical noncanonical 6mer seed and either different (red circles) or identical sequences (blue circles) in the supplementary pairing region position 13-16. p-values were calculated using Pearson correlation analysis.

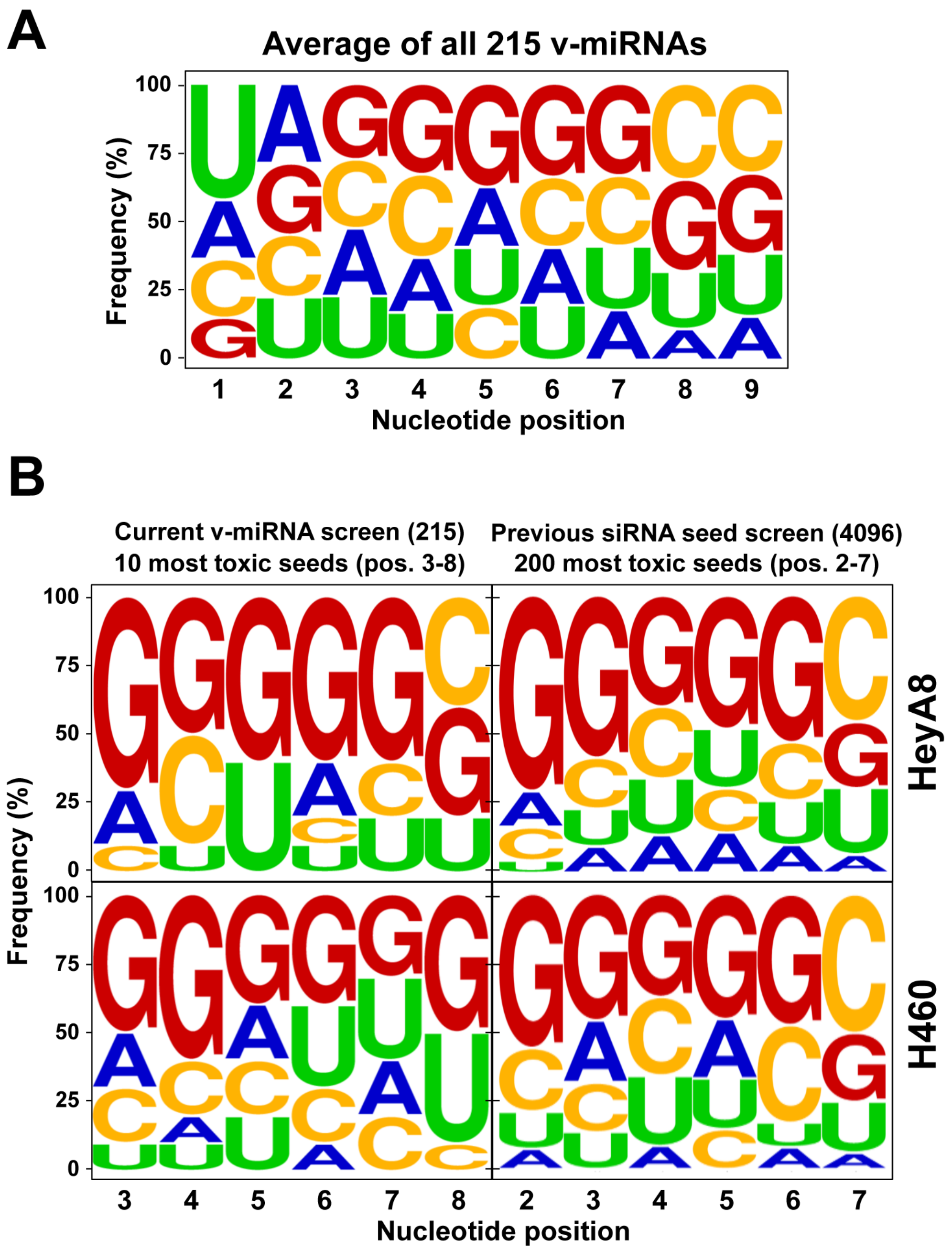


**Figure S2: Related to Figure 4. The composition of the noncanonical 6mer seed of the most toxic v-miRNAs is very similar to the most toxic 2-7 6mer seeds.**

(A) Average nucleotide composition at each of the first 9 nucleotide positions in all 215 v-miRNAs. (B) Average nucleotide composition at each of the 6 seed positions of the noncanonical 6mer seed in the ten most toxic v-miRNAs and the 200 most toxic seeds for the two human cell lines in the 4096 seed siRNAs screen (Gao et al., 2018).

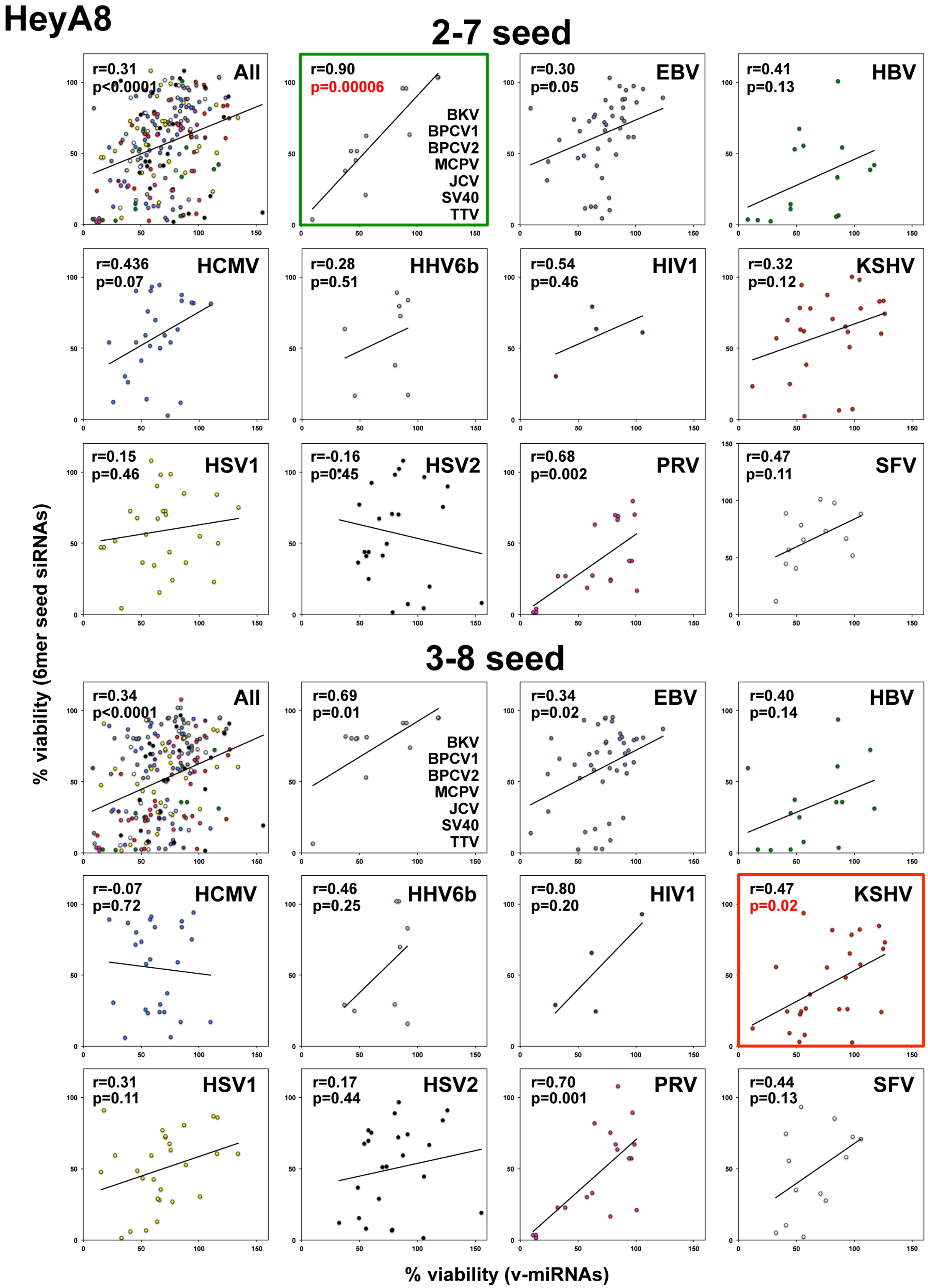
HeyA8
2-7 seed
r=0.31 p<0.0001 All
r=0.90 p=0.00006 BKV BPCV1 BPCV2 MCPV JCV SV40 TTV
r=0.30 p=0.05 EBV
r=0.41 p=0.13 HBV
r=0.436 p=0.07 HCMV
r=0.28 p=0.51 HHV6b
r=0.54 p=0.46 HIV1
r=0.32 p=0.12 KSHV
r=0.15 p=0.46 HSV1
r=-0.16 p=0.45 HSV2
r=0.68 p=0.002 PRV
r=0.47 p=0.11 SFV
3-8 seed
r=0.34 p<0.0001 All
r=0.69 p=0.01 BKV BPCV1 BPCV2 MCPV JCV SV40 TTV
r=0.34 p=0.02 EBV
r=0.40 p=0.14 HBV
r=-0.07 p=0.72 HCMV
r=0.46 p=0.25 HHV6b
r=0.80 p=0.20 HIV1
r=0.47 p=0.02 KSHV
r=0.31 p=0.11 HSV1
r=0.17 p=0.44 HSV2
r=0.70 p=0.001 PRV
r=0.44 p=0.13 SFV
% viability (6mer seed siRNAs)
% viability (v-miRNAs)

**Figure S3. Related to Figure 4. Correlation between toxicity exerted by v-miRNAs in HeyA8 cells and their 6mer seeds in individual viruses.**

Regression analysis showing correlation between the toxicity observed in HeyA8 cells 96 hours after transfection with either 25 nM of the 215 v-miRNAs, or 10 nM of the siRNAs carrying matching 6mer seeds (position 2-7, top) or 10 nM of the siRNAs carrying matching 6mer seeds (position 3-8, bottom) as tested previously (Gao et al., 2018). The data are broken down into individual viruses that code for at least 4 different miRNAs. The viruses with less than 4 miRNAs were combined into one analysis (second plot). Green box: viruses that show a significantly better correlation when the 2-7 6mer seed are considered. Red box: viruses that show a significantly better correlation when the 3-8 6mer seed are considered. p-values were calculated using Pearson correlation analysis. The viruses that are part of the analysis are: BK virus (BKV), bandicoot papillomatosis carcinomatosis virus type 1 and 2 (BPCV1/2), John Cunningham virus (JCV), simian virus 40 (SV40), Torque teno virus (TTV), Epstein-barr virus (EBV), hepatitis B virus (HBV), human cytomegalovirus (HCMV), human herpes virus 6b (HHV6b), human immune deficiency virus 1 (HIV-1), Kaposi sarcoma associated herpes virus (KSHV), Merkel cell polyomavirus (MCPV), herpes simplex virus 1 and 2 (HSV1/2), pseudorabies virus (PRV), and Semliki Forest virus (SFV).

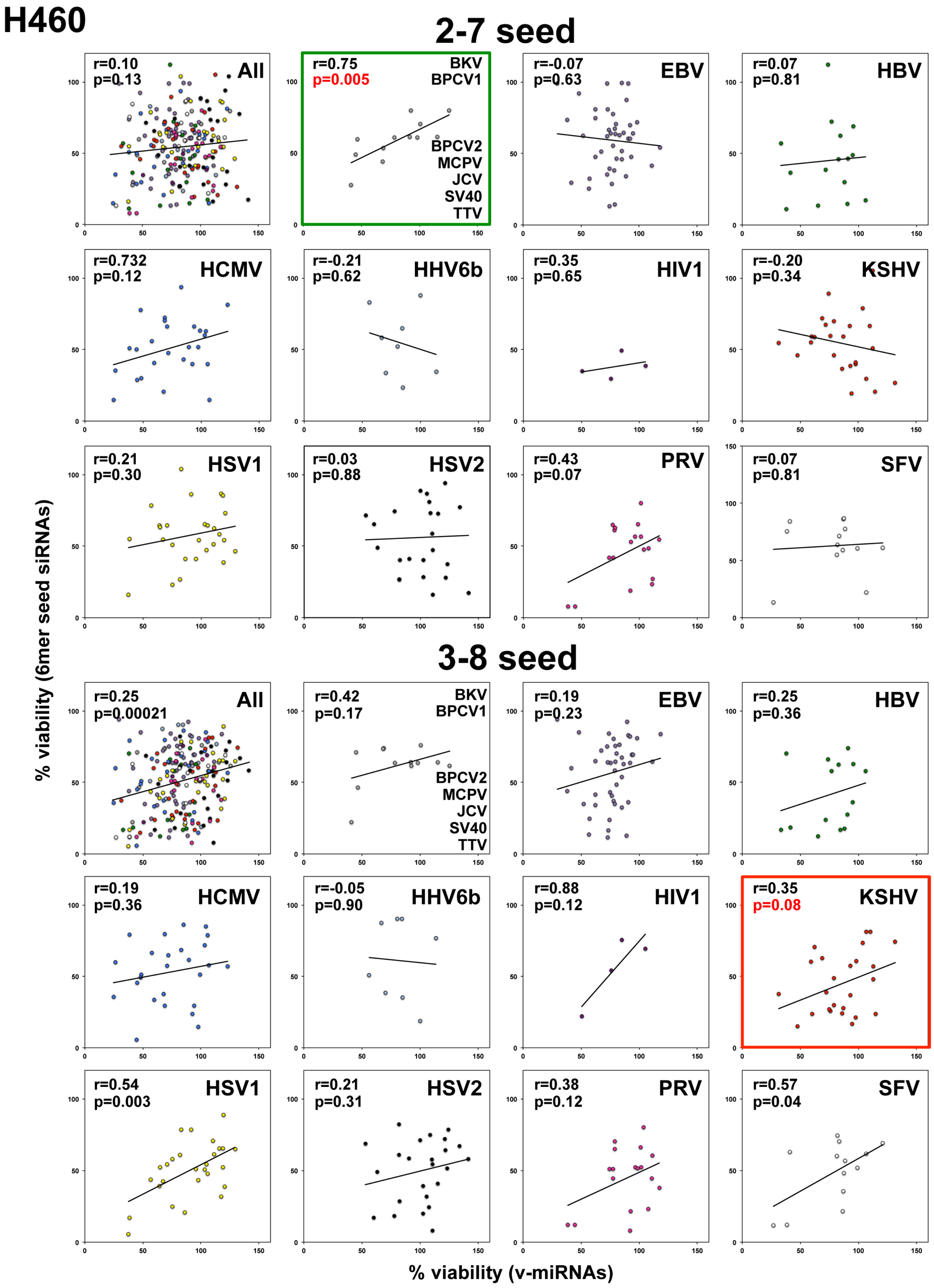
H460
2-7 seed
r=0.10
p=0.13
All
r=0.75
p=0.005
BKV
BPCV1
BPCV2
MCPV
JCV
SV40
TTV
r=-0.07
p=0.63
EBV
r=0.07
p=0.81
HBV
r=0.732
p=0.12
HCMV
r=-0.21
p=0.62
HHV6b
r=0.35
p=0.65
HIV1
r=-0.20
p=0.34
KSHV
r=0.21
p=0.30
HSV1
r=0.03
p=0.88
HSV2
r=0.43
p=0.07
PRV
r=0.07
p=0.81
SFV
3-8 seed
r=0.25
p=0.00021
All
r=0.42
p=0.17
BKV
BPCV1
BPCV2
MCPV
JCV
SV40
TTV
r=0.19
p=0.23
EBV
r=0.25
p=0.36
HBV
r=0.19
p=0.36
HCMV
r=-0.05
p=0.90
HHV6b
r=0.88
p=0.12
HIV1
r=0.35
p=0.08
KSHV
r=0.54
p=0.003
HSV1
r=0.21
p=0.31
HSV2
r=0.38
p=0.12
PRV
r=0.57
p=0.04
SFV
% viability (6mer seed siRNAs)
% viability (v-miRNAs)

**Figure S4. Related to Figure 4. Correlation between toxicity exerted by v-miRNAs in H460 cells and their 6mer seeds in individual viruses.**

Regression analysis showing the correlation between the toxicity observed in H460 cells 96 hours after transfection with either 25 nM of the 215 v-miRNAs or 10 nM of the siRNAs carrying matching 6mer seeds (position 2-7, top) or 10 nM of the siRNAs carrying matching 6mer seeds (position 3-8, bottom) as tested previously (Gao et al., 2018). The data are broken down into individual viruses that code for at least 4 different miRNAs. The viruses with less than 4 miRNAs were combined into one analysis (second plot). Green box: viruses that show a significantly better correlation when the 2-7 6mer seed is considered. Red box: viruses that show a significantly better correlation when the 3-8 6mer seed is considered. p-values were calculated using Pearson correlation analysis. For virus abbreviations see **Figure S3**.

**Table S1: List of all miRNAs encoded by human pathogenic viruses according to miRbase v. 22.1**

**Table S2: Results of toxicity screen of 215 v-miRNAs.**

**Table S3: The 196 6mer seeds that are represented by the 215 v-miRNAs tested.**

**Table S4: Results of screen of the 196 6mer seeds found in the 215 v-miRNAs in a neutral siRNA backbone.** The current screen was performed at 25 nM. The results for the same siRNAs from the previous screen (at 10 nM) are shown for comparison.

**Table S5: Results of the screen of the 197 noncanonical 6mer seeds found in the 215 v-miRNAs.**

**Movie S1: Time lapse video of HeyA8 cells transfected with 10 nM of miR-Ctr1.**

**Movie S2: Time lapse video of HeyA8 cells transfected with 10 nM of hsa-miR-16-5p.**

**Movie S3: Time lapse video of HeyA8 cells transfected with 10 nM of kshv-miR-K12-6-5p.**
